# A natural antisense to brain-derived neurotrophic factor impairs extinction of drug seeking

**DOI:** 10.1101/851949

**Authors:** Neil A. Youngson, Matthew R. Castino, Angela Stuart, Kelly A. Kershaw, Nathan M. Holmes, Paul D. Waters, Kevin V. Morris, Kelly J. Clemens

**Affiliations:** School of Medical Sciences, University of New South Wales, Sydney, Australia; School of Psychology, University of New South Wales, Sydney, Australia; School of Biotechnology and Biomolecular Sciences, University of New South Wales, Sydney, Australia; Center for Gene and Cell Therapy, Beckman Research Institute at the City of Hope, CA, USA

**Author notes:** Corresponding Author: Kelly J. Clemens, School of Psychology, University of New South Wales, UNSW Sydney, NSW 2052, Australia.

**Keywords:** Nicotine, addiction, extinction, BDNF, non-coding RNA, epigenetics

## Abstract

**BACKGROUND:** Brain derived neurotrophic factor (BDNF) is critical for the extinction of drug-seeking. Expression of the *Bdnf* gene is highly regulated via interactions with non-coding RNA, which themselves are altered following drug exposure. Here we investigate whether a novel long non-coding RNA antisense to *Bdnf* prevents extinction of drug-seeking. METHODS: Strand-specific RNA sequencing identified a novel long non-coding RNA antisense to exon IV of the *Bdnf* gene in the ventromedial prefrontal cortex of 8 adult male rats. We then assessed *asBdnf-IV* expression using strand-specific reverse transcription and quantitative polymerase chain reaction following acquisition, extinction or abstinence from intravenous nicotine self-administration (N = 116). A functional role of the *asBdnf-IV* in extinction of nicotine-seeking was established by infusing gapmer oligonucleotides into the infralimbic cortex prior to extinction and testing for the effect of these infusions on reinstatement and reacquisition of nicotine-seeking (N = 36).

**RESULTS:** RNA sequencing identified the presence of a novel long non-coding RNA antisense to exon IV of the *Bdnf* gene (*asBdnf-IV*). Expression of *asBdnf-IV* was elevated following intravenous nicotine self-administration but not experimenter-administered nicotine. Elevated *asBdnf-IV* persisted across abstinence and to a greater extent following extinction training, suggesting an interaction between abstinence and extinction learning. In support of this, knockdown of the *asBdnf-IV* across extinction, but not abstinence, significantly attenuated nicotine-primed reinstatement of nicotine-seeking.

**CONCLUSIONS:** *asBdnf-IV* accumulates in the infralimbic cortex across self-administration training, interferes with the inhibitory learning that underpins extinction of drug-seeking, and predisposes animals to drug relapse.

---

Tobacco dependence is a significant health burden worldwide, killing more than 5 million people each year (1). Public health initiatives have helped to reduce the initiation of smoking, but have done little to encourage abstinence among established smokers (2). Abstinence requires development of inhibitory control over cravings that are triggered by smoking-related cues, and precipitate relapse (3). Neuroimaging studies have shown that the development and maintenance of this control (in the form of dampened cue-reactivity) correlates with neuronal activity in the prefrontal cortex (4, 5), particularly its ventromedial portion (vmPFC) (6). However, at present, very little is known about the cellular and molecular processes within this region that underlie the development and maintenance of inhibitory control over drug-seeking.

Research in rodents has shown that the vmPFC plays a critical role in regulating relapse to drug-seeking (7). Specifically, studies of drug self-administration in rats have shown that extinction of established drug-seeking requires neuronal activity in the infralimbic cortex (IL; rodent homologue of the vmPFC) (8), including expression of the immediate early gene, brain derived neurotrophic factor (*Bdnf*). BDNF infused into the IL enhances the extinction of cocaine conditioned place preference (9), and significantly attenuates reinstatement of cocaine-seeking behaviour (10). The implication of these findings is that BDNF signalling in the IL is important for the acquisition and maintenance of inhibitory memories that form during extinction of drug-seeking behaviour. These inhibitory memories oppose or mask the tendency towards drug-seeking, and thereby, underlie successful abstinence from drug-seeking behaviour.

Recent work in our laboratory has shown that, within the IL, extinction of nicotine self-administration is associated with epigenetic modifications in the promoter region of *Bdnf* exon IV, an activity-dependent *Bdnf* splice variant that is increased following exposure to drugs of abuse (11). These changes include decreases in the repressive epigenetic marks H3K27me^3^ and H3K9me^2^, thereby promoting a permissive chromatin state (12). Importantly, despite these changes, *Bdnf-IV* mRNA expression was unchanged, suggesting that nicotine self-administration epigenetically primes *Bdnf-IV* expression, and that this priming persists across extinction in the absence of detectable changes in gene expression. The processes that implement and maintain these changes are not yet known; however, recent work suggests that non-coding RNAs may play a critical role (13). Specifically, an antisense *Bdnf* (*asBdnf)* opposite exon IX of the *Bdnf* gene has recently been shown to recruit elements of a repressive complex to the H3K27me^3^ mark, effectively silencing *Bdnf-IX* mRNA transcription (14). Accordingly, we hypothesized that, within the IL, an *asBDNF* may regulate nicotine-induced modifications to histone H3 in the promoter region of *Bdnf* exon IV; and that by targeting this antisense transcript, we might influence the extinction and reinstatement of nicotine-seeking behaviour.

The present study tested these hypotheses. We first discovered a new natural transcript antisense to exon IV of the *Bdnf* gene (*asBdnf-IV*) in the rat IL. We then showed that self-administration of nicotine leads to an upregulation of *Bdnf-IV* mRNA and *asBdnf-IV* in this region (changes were not evident following passive nicotine injections), and that the level of *asBdnf-IV* further increased following extinction compared to abstinence. Finally, we confirmed a functional relationship between *asBdnf-IV* in the IL and nicotine-seeking behaviour, as knockdown of the transcript across extinction of nicotine self-administration (but not abstinence) significantly attenuated reinstatement of nicotine-seeking.

## METHODS and MATERIALS

### Animals and Housing

Male Sprague Dawley rats (175-200 g; Australian Resources Centre, Perth, Australia) were housed, four per cage with a 12 h reverse dark/light cycle (light on at 19:00 hours, off at 07:00 hours). Rats received *ad libitum* access to food and water until two days prior to nicotine self-administration when food was reduced to 22 g/rat/day. All testing was carried out during the dark cycle.

All procedures were approved by the Animal Care and Ethics Committee of the University of New South Wales and were conducted in accordance with the Australian Code for the Care and Use of Animals for Scientific Purposes (8^th^ ed.).

### Drugs

Nicotine tartrate was dissolved in sterile saline (0.9% NaCl) and administered intravenously at 30 μg/kg/100 μL infusion or intraperitoneally at 0.3 mg/kg. All concentrations of nicotine refer to the base and solutions were pH adjusted to pH 7.4 using 1M sodium hydroxide.

### Tissue Processing and RNA Extraction

Rats are euthanised with pentobarbitone sodium (325 mg/ml/rat), brains removed and tissue either immediately frozen in liquid nitrogen for later micropunching, or the ventromedial prefrontal cortex (vmPFC; primarily containing the infralimbic cortex, 2 mm^2^; bregma: 5.16 mm – 2.76 mm) was dissected onto dry ice using a scalpel blade. Micropunches were taken from 1 mm sections and the region of interest punched with a 1 or 2 mm disposable micropunch according to the rat brain atlas (15). RNA was then isolated using the Trizol extraction method according to manufacturer’s instructions (Invitrogen, CA, USA).

### RNA Sequencing

RNA quality was assessed using the Agilent Bioanalyzer (Agilent, CA, USA), with all samples returning RIN numbers between 8.3 and 8.8. Total RNA was ribosome depleted in preparation for Illumina TruSeq stranded RNA library construction and sequencing at the Ramaciotti Centre for Genomics (UNSW, Australia). Approximately 25 million 100bp paired-end reads were generated for each sample, which were mapped to the rat genome using hisat2 (16) (with the options: known-splice site-in file Rattus_norvegicus.Rnor_6.0.84.txt; rna-strandness RF). Read counts were conducted with htseq-count (17) (with the options -m union, -s yes). Normalized and relative transcript levels were calculated with edgeR (18).

### Validation of Gene Target

An RNA transcript antisense to exon IV of the *Bdnf* gene was selected for further investigation due to associations between the *Bdnf-IV* splice variant and both addiction and extinction learning (11). We designed a number of primers to initially validate the transcript antisense to *Bdnf* exon IV in the mPFC using strand-specific reverse transcription followed by quantitative PCR (RT-qPCR).

Total RNA was DNase-treated (Sigma DNAse I) and reverse transcribed in a C1000 Touch Thermal Cycler (Bio-Rad Laboratories, CA, USA) using the Invitrogen SuperScript III First-Strand Synthesis System for RT-PCR (Thermofisher) according to the manufacturer’s instructions. Quantitative PCR was performed in a StepOnePlus system with the use of SYBR Select and PowerUP SYBR Master Mix (Applied Biosystems, VIC, Australia). All qPCR primers (Integrated DNA Technologies, IA, USA) were designed using Primer3 software (19) and verified using the BLAST-like alignment tool (BLAT; (20). Primer efficiencies were determined for each primer pair using a standard curve, before specificity of target sequences was confirmed with 1.5% agarose gels and Sanger Sequencing (Ramaciotti Centre for Genomics, UNSW). The primer pair giving the most reliable amplification was selected for future studies. For each target (run in triplicate for each sample) RNA levels were normalized to the housekeeping genes GAPDH and β-actin.

### Experimenter Administered Nicotine

To examine *asBdnf-IV* expression in response to experimenter-administered nicotine, 48 rats received either 12 injections of saline (SAL), 11 injections of saline and then one of nicotine (ACUTE NIC) or 12 injections of nicotine (CHRONIC NIC) on 12 consecutive days. Either 2 or 24 hours after the final injection, rats were euthanised and tissue processed for gene expression as described above.

### Intravenous Catheter Surgery and Intravenous Self-Administration

Rats underwent surgery for the implantation of a chronic intravenous catheter as described previously (21). Following recovery from surgery, behavioural training began with two daily one hr habituation sessions conducted in standard self-administration equipment (Med Associates, VT, USA; (21)) with nose-pokes covered and the house-light on. Rats were then trained to self-administer nicotine (Nic) or saline (Sal) for 12 days on a fixed ratio-1 (FR-1) schedule of reinforcement. Self-administration sessions lasted for one hr and began with the illumination of the house-light. Each response on the active nose-poke resulted in a single infusion of nicotine (30 µg/kg/100 µL; 3 s) or saline (100 µL over 3 s), presentation of the cue-light inside the nose-poke (3 s) and termination of the house-light (20 s). Responses during the 20 s time-out period or on the inactive nose-poke were recorded but were of no scheduled consequence. Extinction training involved removal of nicotine and the associated cue for 1 or 6 days. Responses on the active nose-poke were recorded but were of no scheduled consequence. Rats allocated to the abstinence condition were handled daily but did not enter the test room or chambers.

Thirty minutes after the end of the session (90 minutes after the beginning of the session) or at an equivalent time on abstinence days, rats were anaesthetised and tissue processed as described above.

### Intracranial Surgery and Microinjection Procedures

For knockdown studies, rats were stereotaxically implanted with a 6 mm 26-gauge bilateral cannula targeting the infralimbic cortex (AP 3.0, ML 0.6, DV 3.5 mm) immediately after implantation of the IV catheter (21). Rats underwent training as described above, including three days of extinction training. This was followed by a single reinstatement test where a single injection of nicotine (0.3 mg/kg s.c.) was administered immediately prior to a 90 minute extinction session where cue-lights were presented, but nicotine was not.

Oligonucleotides were designed based on regions of interest selected from the RNA-seq data and submitted to Integrated DNA Technologies (IDT, Singapore) for validation and design of gapmer oligonucleotides. Two synthetic gapmer oligonucleotides 20 nucleotides in length containing 2-O’-methyl RNA molecules and directed against the *asBdnf-IV* or scrambled control were selected to be highly specific to RNA (14). The OGNs targeting the *asBdnf-IV* were combined for each infusion and compared with a scrambled control. Oligonucleotides (2nM/μL, 1 μL/side) were infused at a rate of 0.25 μL/min into the infralimbic cortex 90 mins prior to the first extinction session only, or at an equivalent time of day for rats in the abstinence condition (22). Infusion cannula extended 1 mm below the guide cannula and remained in place for 2 minutes post-infusion to allow diffusion of the solution. Rats with blocked cannula or damage to the pedestal received sham infusions where rats were restrained but only received unilateral or no infusion. No differences between these animals and those successfully receiving the scrambled control infusion were detected and therefore data was combined.

To validate knockdown and verify cannula placement rats received a second infusion of the OGNs or scrambled control a minimum of three weeks after the final self-administration session. Treatment allocation was counterbalanced based on previous infusion. After the infusion, rats were returned to the home cage. Three days later, they were euthanized and their brains removed, frozen in liquid nitrogen, and stored at −80°C. For analysis of cannula placement and verification of knockdown, the prefrontal cortex was sliced into 1 mm sections using a rat brain matrix. Slices were photographed for later verification of cannula placement. A 2 mm micropunch was then taken from each hemisphere, centered around the tip of the cannula tract in the infralimbic cortex. Punches were combined into a single tube and then stored at −80°C until processing for gene expression as described above.

### Statistical Analysis

All data were analysed using a mixed model ANOVA, repeated measures ANOVA or t-tests as appropriate. For qPCR data, when the Ct SD of triplicates was greater than 0.5, the outlier was identified and excluded, or if necessary, the sample was rerun.

## RESULTS

### Identification of rat *asBdnf-IV* transcript

Strand-specific RNA-Seq identified the presence of transcripts antisense to the *Bdnf* gene (Fig. 1a,b). Using the RNA-seq data, and because of past associations between addiction and the Exon IV splice variant of *Bdnf* (11) we selected a transcript antisense to exon IV of the *Bdnf* gene for further investigation.

**Figure 1.**
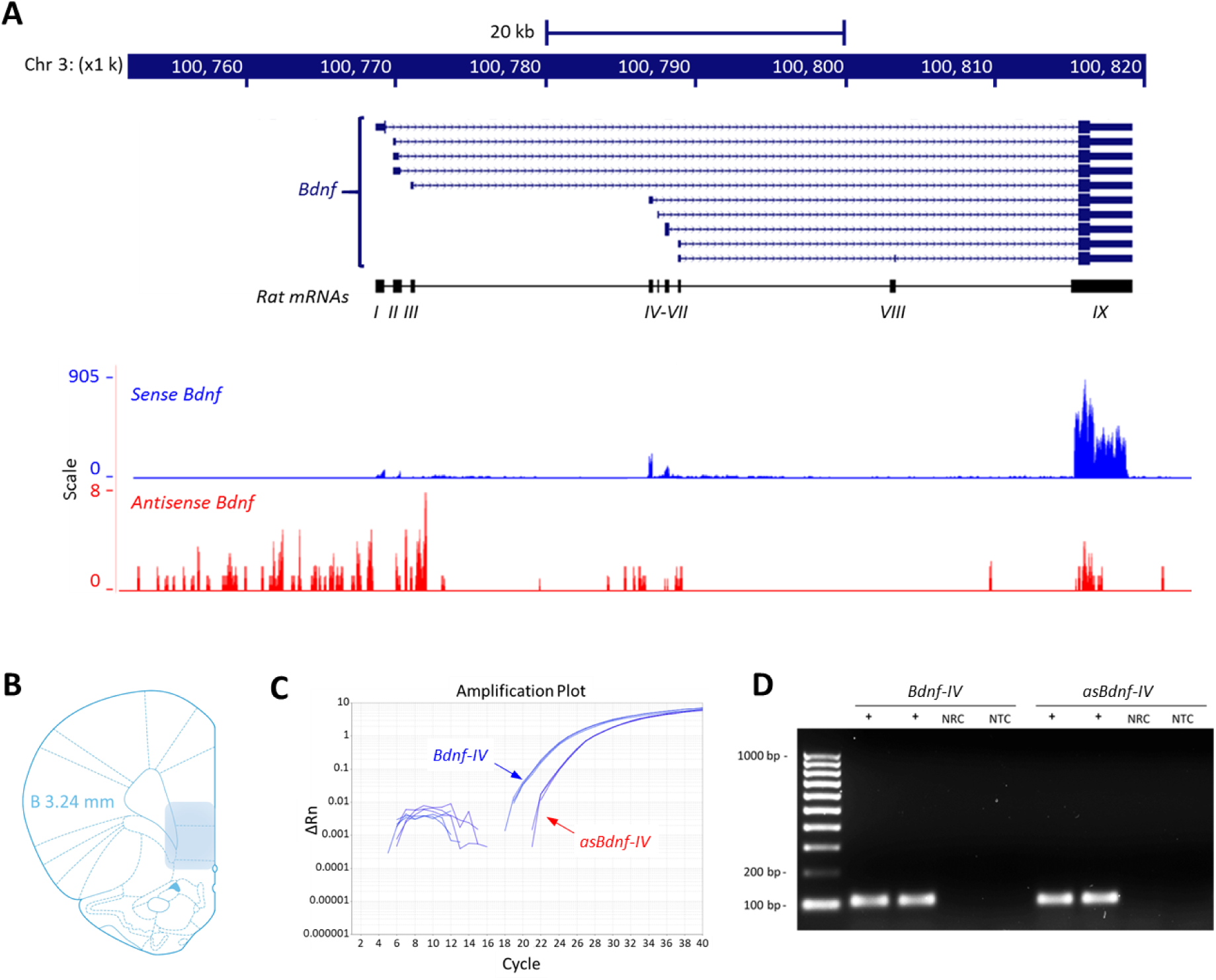
Characterization of the asBdnf-IV transcript. **A)** RNA-Seq data confirming the presence of an antisense Bdnf (asBdnf; red) transcript to the Bdnf gene (blue) in the prefrontal cortex of naïve rats (chr 3: 100,751,440 - 100,823,456 [72,017 bp], adapted from UCSC browser (20)). **B)** Area of tissue dissected for all analyses; **C)** Amplification plot and **D)** gel showing PCR-product following strand-specific reverse transcription and PCR; NRC: no reverse transcriptase control; NTC: no template control.

### Expression of *Bdnf-IV* and *asBdnf-IV* in the rat brain

The expression of *Bdnf* mRNA in the rat brain has been well documented (23), but the distribution of the *asBdnf-IV* is unknown. We measured the levels of *Bdnf-IV* and *asBdnf-IV* transcripts across five regions of interest in the brain of male Sprague-Dawley rats (Fig. 2A). Both transcripts were abundant in the medial prefrontal cortex and hippocampus (Fig. 2B). In particular, high levels of expression were evident in the IL and PL, brain regions that have been implicated in extinction and reinstatement of drug-seeking (8). Consistent with previous findings in the Rhesus monkey (14), the *Bdnf-IV* mRNA levels were approximately 9-15 times more abundant than *asBdnf-IV* and varied across brain regions (Fig. 2C).

**Figure 2.**
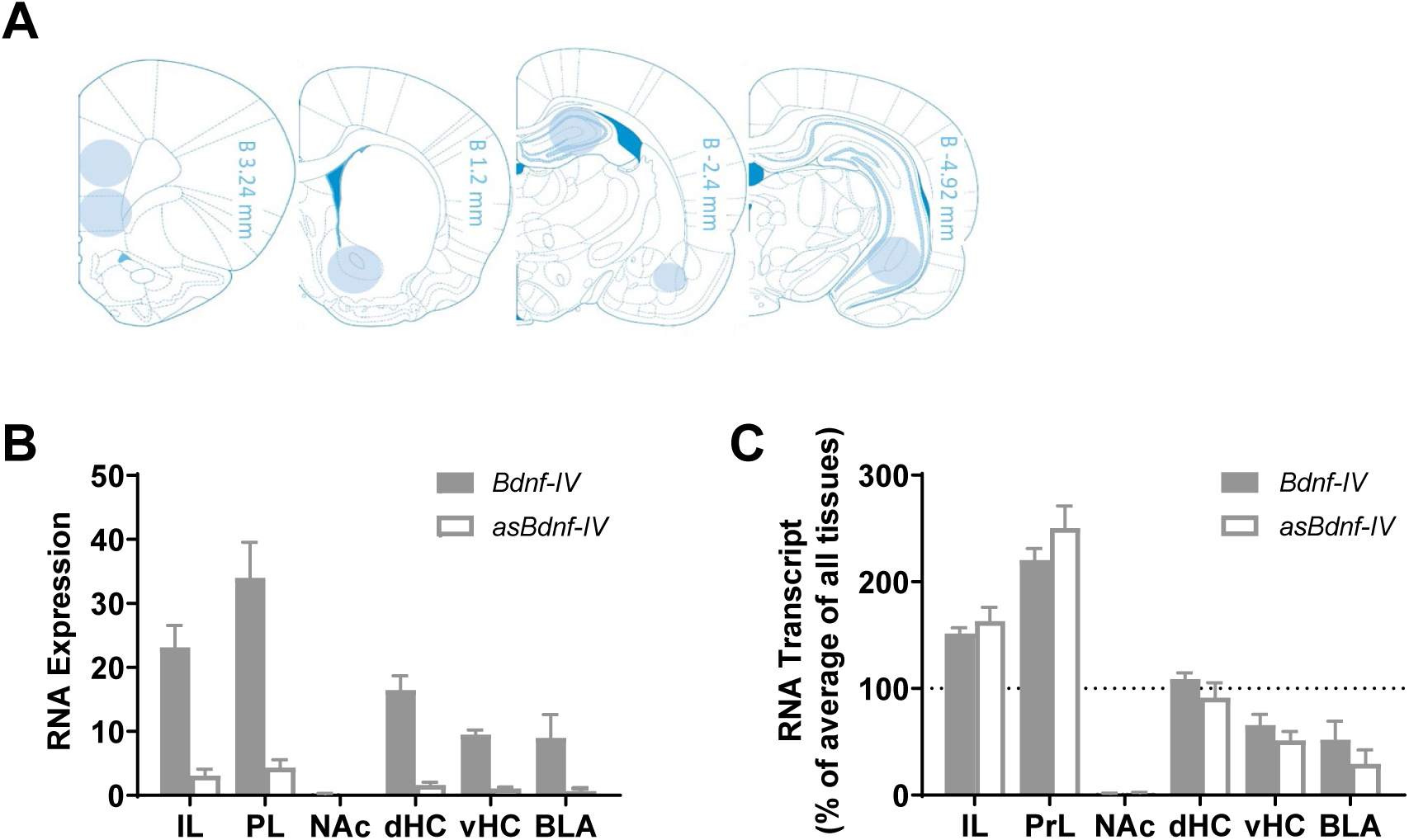
Distribution of Bdnf-IV mRNA and asBdnf-IV across the rat brain. **A)** Brain regions punched for analysis with distance from Bregma indicated. **B)** Comparison of expression levels and **C)** relative expression as a percentage of average expression across all tissues. IL: infralimbic cortex; PrL: prelimbic cortex; NAc: nucleus accumbens; dHC: dorsal hippocampus; vHC: ventral hippocampus; BLA: basolateral amygdala. n = 4.

### *asBdnf-IV* is not altered following systemic injection of nicotine

To determine if *asBdnf-IV* is altered following passive administration of nicotine, naïve rats were administered acute or chronic (12 days) injections of nicotine. Irrespective of drug administration, the injection procedure produced a significant increase in both *Bdnf-IV* and *asBdnf-IV* expression 2-hrs post-injection that returned to baseline 24 hours later (Fig. 3). This result suggests that neither acute nor chronic experimenter-administered nicotine produces a sustained increase in *asBdnf-IV* expression.

**Figure 3.**
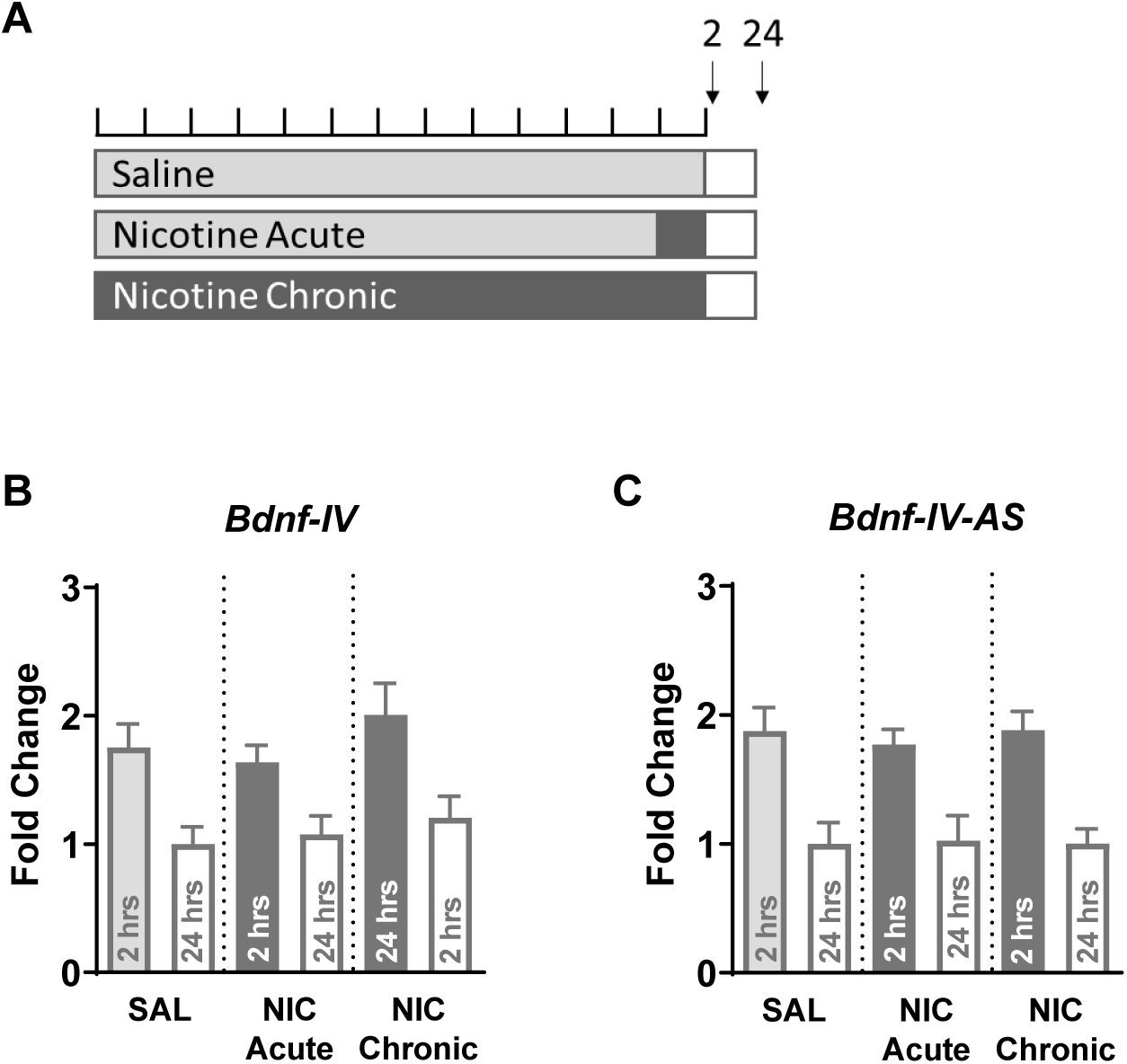
Bdnf-IV mRNA and asBdnf-IV expression following 12 daily injections of saline, 11 days of saline and 1 of nicotine (Nicotine acute) or 12 days of nicotine (Nicotine chronic). **A)** Experimental design. Rats receive 12 daily injections of saline (SAL), 11 daily injections of saline followed by a single injection of nicotine (NIC ACUTE) or 12 daily injections of nicotine (NIC CHRONIC) and euthanized 2 or 24 hours after the final injection. **B)** Bdnf-IV mRNA expression significantly increased following the injection procedure (main effect of time: F(1,42)=24.79, p<0.001) irrespective of treatment (F<1). **C)** asBdnf-IV expression significantly increased following the injection procedure (main effect of time: F(1,38)=42.38, p<0.001) irrespective of treatment (F<1). N=7-8 rats per group.

### *asBdnf-IV* is increased following intravenous nicotine self-administration

To explore the relationship between nicotine self-administration and the expression of both *Bdnf-IV* mRNA and *asBdnf-IV* we assessed the expression of these transcripts in rats that had self-administered nicotine. Compared to self-administration of saline, nicotine self-administration produced a significant increase in both *Bdnf-IV* mRNA and *asBdnf-IV* that persisted across one day of extinction training (Fig. 4B, C). *asBdnf-IV*, but not *Bdnf-IV*, expression was positively correlated with total active responses across training (Fig. 4D, E), suggesting that both *Bdnf-IV* mRNA and *asBdnf-IV* increase as a consequence of active self-administration, and that this increase persists in the absence of nicotine intake.

**Figure 4.**
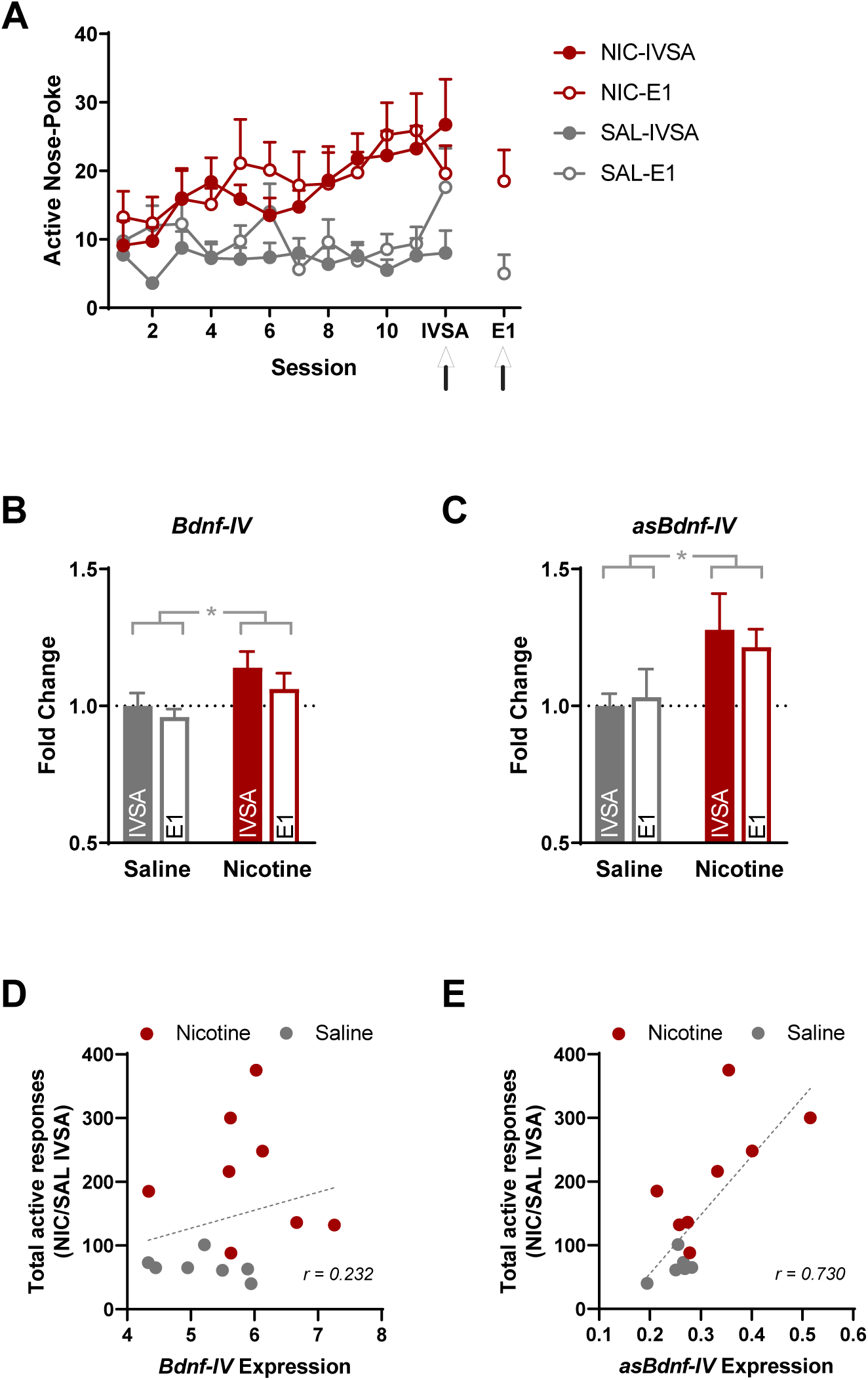
Bdnf-IV-mRNA and asBdnf-IV expression following nicotine self-administration and brief extinction. **A)** Rats self-administering nicotine made significantly more active nose-pokes than rats responding for saline (main effect of drug (F(1,25)=7.110, p=0.013, 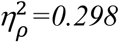), although there were no differences in acquisition between rats allocated to the IVSA versus EXT test conditions (F=0.034). Arrows indicate the two time points (90 min post-session) when tissue was collected. **B)** Nicotine self-administration resulted in a significant increase in Bdnf-IV sense mRNA (main effect of drug: F(1,25)=5.744, p=0.024, 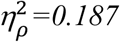), that did not differ between the last day of self-administration and the first day of extinction (man effect of test: F=1.374; interaction F=0.139). **C)** Nicotine self-administration resulted in a significant increase in asBdnf-IV transcript (main effect of drug: F(1,25)=5.350, p=0.029, 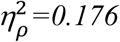), that did not differ between the last day of self-administration and the first day of extinction (main effect of test: F=0.034; interaction F=0.264). **D)** Bdnf-IV expression and total active responses across training in the IVSA group were not correlated (r(15)=0.232, p=0.85), however **E)** asBdnf-IV and total active responses in the IVSA group were highly correlated (r(15)=0.730, p=0.002). N=6-8/group, asterisks indicate significant difference when p<0.05. N =8/group.

### Increased *asBdnf-IV* persists across abstinence

To examine the persistence of elevated *asBdnf-IV* across extinction we assessed changes in gene expression following six days of extinction training compared to those that occur simply as a consequence of prior nicotine exposure (abstinence). This is the same time point at which we have previously observed decreases in H3K27 trimethylation and H3K9 dimethylation at the BDNF exon IV promoter (12). Rats underwent 12 days of self-administration followed by 6 days of extinction training (Fig. 5A) and euthanasia (E6), or 6 days of forced abstinence in the home cage and euthanasia (ABS). Rats in both the saline and the nicotine infusion groups showed a significant increase in *Bdnf-IV* mRNA following extinction training compared to rats in the abstinence groups (Fig. 5B). This indicates that *Bdnf-IV* mRNA is increased in response to extinction training independently of prior nicotine history. In contrast, *asBdnf-IV* expression was greater in those rats that had previously self-administered nicotine compared to saline controls, independently of whether they had been through extinction or abstinence; and was greater in those rats that had been through extinction rather than just abstinence, independently of whether they had previously self-administered nicotine or saline. Together these findings suggest that nicotine self-administration and extinction have additive effects on *asBdnf-IV* expression.

**Figure 5.**
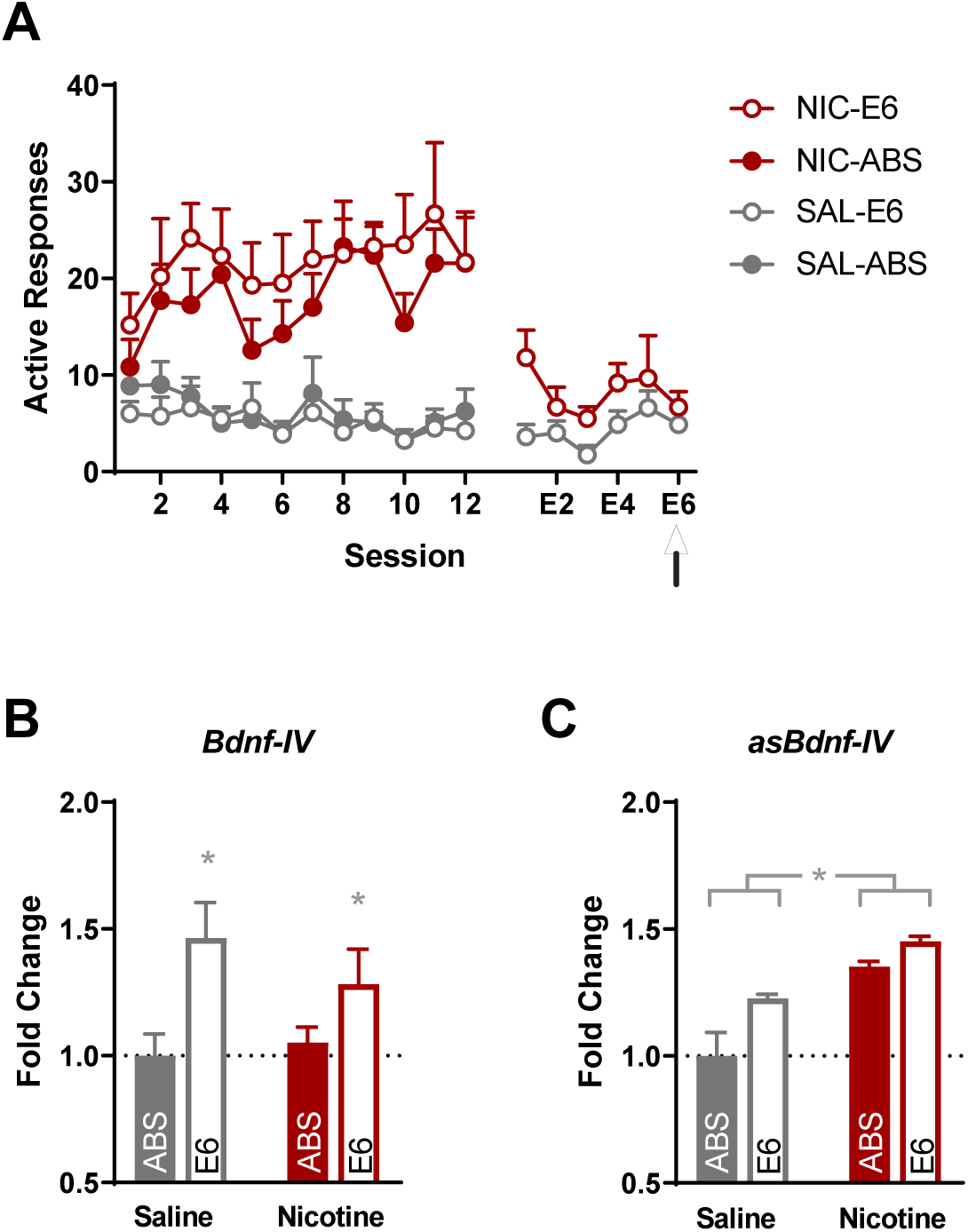
Bdnf-IV mRNA and asBdnf-IV expression following nicotine self-administration followed by 6 days of abstinence or extinction. **A)** Rats self-administering nicotine made significantly more active nose-pokes than rats responding for saline (main effect of drug (F(1,24)=46.149, p<0.001, 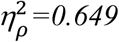), although there were no differences in acquisition between rats allocated to the EXT6 versus ABS test conditions (F<1). **B)** Extinction training following self-administration resulted in a significant increase in Bdnf-IV Sense mRNA (main effect of extinction condition: F(1,24)=8.830, p=0.007, 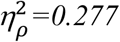), independently of self-administration infusion solution (F<1). **C)** Nicotine self-administration resulted in a significant increase in asBdnf-IV transcript (main effect of drug: F(1,24)=5.016, p=0.035, 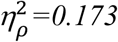), that did not differ between animals receiving extinction versus abstinence (F<1). N=6-8/group.

### Knockdown of *asBdnf-IV* in the IL across extinction training attenuates relapse

To demonstrate a causal relationship between *asBdnf-IV* expression in the IL and nicotine-seeking behavior, we selectively degraded this transcript using an Anti-*asBdnf-IV* OGN gapmer oligonucleotide (OGN) compared to a scrambled OGN control (SCR). Rats were trained to self-administer nicotine for 12 days. This was followed by either three days of extinction training (EXT) where a response on the nose-poke resulted in presentation of the cue-light, but no infusion; or three days of abstinence (ABS) where rats remained in the home cage. Ninety minutes prior to the first extinction session (EXT group) or at an equivalent time on the first day of abstinence (ABS group) rats were infused with a combination of two Anti-*asBdnf-IV* OGNs or scrambled control oligonucleotides (Fig. 6A). The OGNs produced a 36% decrease in *asBdnf-IV* expression in tissue punched around the tip of the cannula (Fig. 6B, C).

**Figure 6.**
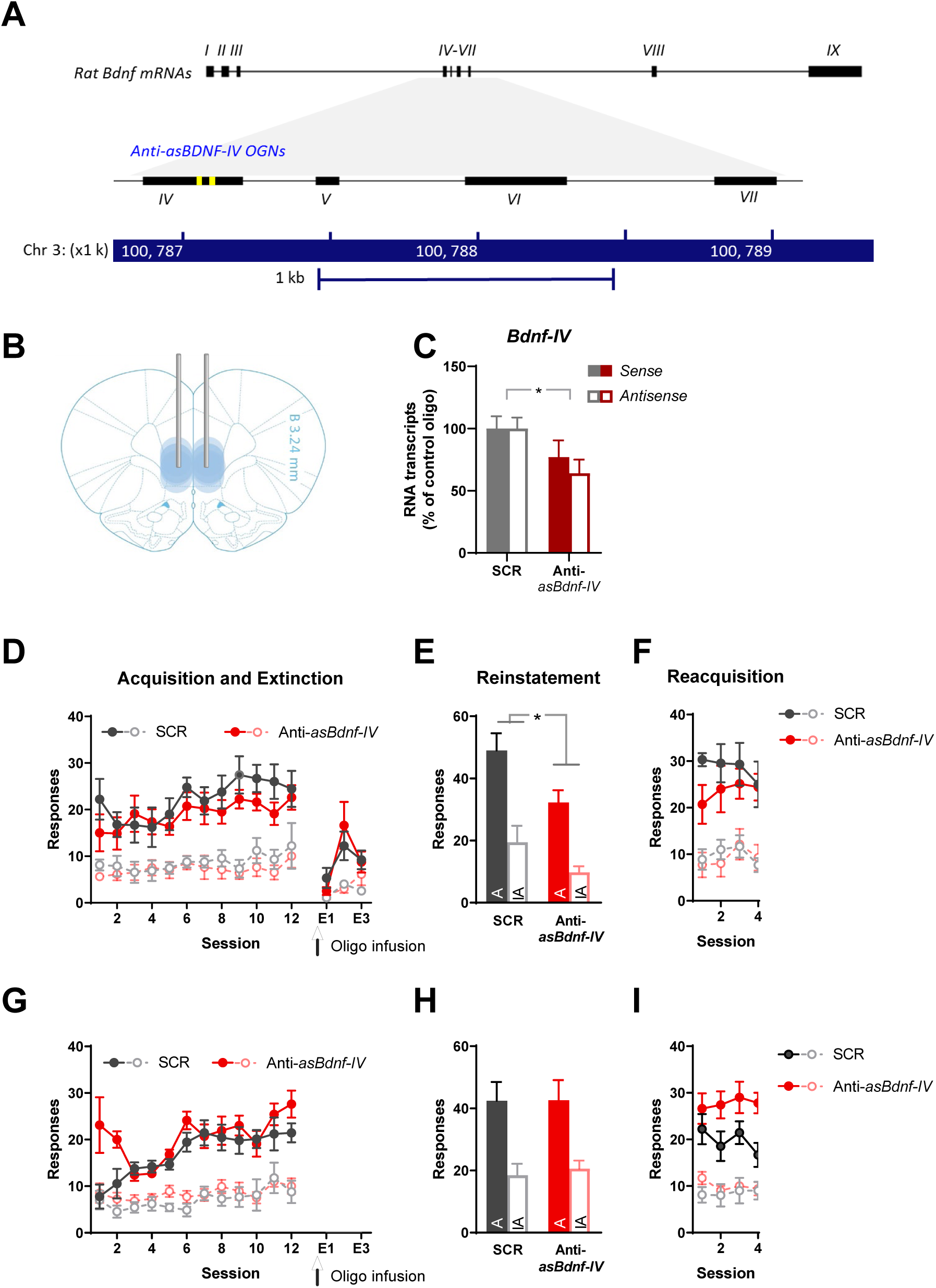
Knockdown of asBdnf-IV significantly attenuated reinstatement of nicotine seeking. **A**) Position of anti-asBdnf-IV oligonucleotides (yellow) in the UCSC Browser (https://genome.ucsc.edu/, Jul. 2014 (RGSC 6.0/rn6) assembly). **B)** Placement of infusion cannula in the infralimbic cortex (IL; all tips within the grey circles) and location of 2mm punches (grey circles) used to verify knockdown. **C)** Verification of asBdnf-IV knockdown by oligonucleotides in the infralimbic (main effect of gene: F(1,8)=28.916, p=0.001; η_p_^2^=0.783; interaction gene by oligo: F(1,8)=7.998, p=0.022; η_p_^2^=0.500 main effect of oligo: F(1,8)=4.328, p=0.071; η_p_^2^=0.351). **D)** Rats rapidly acquired the self-administration response (main effect of day: F(11,165)=2.792, p=0.002, 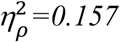; day by nose-poke interaction F(11,165)=1.935, p=0.038; 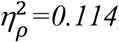) to an equivalent extent in animals subsequently assigned to receive the Anti-asBdnf-IV (Anti-BDNF) or scrambled control (SCR; no effect or interactions involving infusion oligo: Fs<2.11, ps>0.05). No significant differences in responding following the infusion, and across subsequent extinction sessions were observed (No significant effect or interactions involving Anti-BDNF infusion Fs<2). **E)** Infusion with the Anti-asBdnf-IV prior to day 1 of extinction significantly attenuated subsequent relapse to nicotine-seeking (main effect of nose-poke: F(1,15)=15.418, p=0.001; 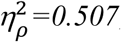; main effect of oligo F(1,15)=8.057, p=0.012; 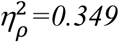). **F)** Anti-asBdnf-IV knockdown did not impact on subsequent reacquisition of self-administration (Fs<1.2). **G)** In a second cohort of rats, all animals rapidly acquired the self-administration response (main effect of day: F(11,187)=2.697, p=0.003, 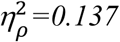; day by nose-poke interaction F(11,187)=2.211, p=0.015; 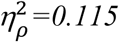). An initial coincident difference between subsequent treatment groups (12 days by OGN interaction (F(11,187)=2.275, p=0.013; 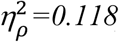), was due to differences across the first 2 days that were no longer evident across the final 10 days of training (10 days by OGN interaction (F(9,153)=1.677, p=0.099; 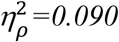). No extinction training was performed in this group. **H)** Infusion with the Anti-asBdnf-IV on day 1 of abstinence had no impact on subsequent relapse to nicotine-seeking (main effect of nose-poke: F(1,17)=20.442, p=0.001; 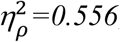; No effect of oligo F(1,17)=0.020, p=0.890; 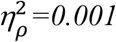). **I)** Anti-asBdnf-IV infusion did not impact on reacquisition, although a decrease in responding by the SCR group was evident (main effect of oligo: F(1,17)=4.748, p=0.04; 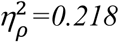). N=8-10 per group.

Infusion with the control OGN had no detectable effect on nose-poke responses across extinction of nicotine-seeking (Fig. 6D). However, knockdown of *asBdnf-IV* in the IL significantly attenuated nicotine-primed reinstatement (Fig. 6E) without impacting on later reacquisition (6F). In contrast, infusion with the Anti-*asBdnf-IV* OGN had no effect on reinstatement after three days of abstinence (group ABS) (Fig. 6G, H, I). Together these results show that the knockdown of *asBdnf-IV* is not due to any effect of the OGN on expression of drug-seeking at the reinstatement test, but rather, is associated with encoding of the extinction memory and its subsequent impact on drug-seeking.

## DISCUSSION

Long non coding RNAs, including natural antisense transcripts, have been associated with a range of diseases and disorders, including neurodegenerative and neuropsychiatric disorders (24). Here we show that a novel natural antisense transcript found opposite exon IV of the *Bdnf* gene maintains a susceptibility to relapse of nicotine-seeking, and does so independently of changes in *Bdnf-IV* mRNA. Together with our previous reports of permissive chromatin in this brain region, our findings indicate *asBdnf-IV* is associated with long-lasting epigenetic changes that persist beyond cessation of drug intake.

Here we provide evidence from multiple sources that nicotine increases transcription of a previously undescribed *asBdnf* in the rat brain antisense to exon IV of the *Bdnf* gene. The relationship between the sense and antisense transcripts is largely concordant, yet dissociable: the antisense transcript is elevated following nicotine self-administration, and in contrast to the sense transcript, remains elevated after either abstinence or extinction training. This finding is consistent with an increasing literature demonstrating the functional diversity in lncRNA modulation of gene expression, with numerous examples of lncRNA promoting a permissive gene environment (25). It also supports the likelihood that multiple asBDNF isoforms exist, and that these may independently influence BDNF gene expression. The potential modulation of Exon IV of the *Bdnf* gene by an antisense transcript is of particular interest given its association with addictive substances (11) and extinction learning (26); and the fact that we have previously detected epigenetic changes in the promoter region of *Bdnf* exon IV (12) using the same behavioural paradigm.

We have previously shown that following 6 days of extinction from nicotine self-administration, H3K27me^3^ is *decreased* in the promoter region of *Bdnf-IV* (12), leading to the hypothesis that the Exon IV antisense transcript induces a permissive chromatin state in the rat IL (Fig. 7). In support of this hypothesis, the Exon IX antisense identified in mice by Modaressi et al (2012) was shown to reduce gene expression via recruitment of repressive epigenetic elements to the H3K27me^3^ mark(14) (Fig 7). The precise nature of the interaction between *asBdnf-IV* and epigenetic elements in the context of nicotine dependence remains to be determined, but it should be further noted that the type of interaction proposed here is consistent with past findings that chronic nicotine leads to an overall relaxed chromatin state (27, 28).

**Figure 7.**
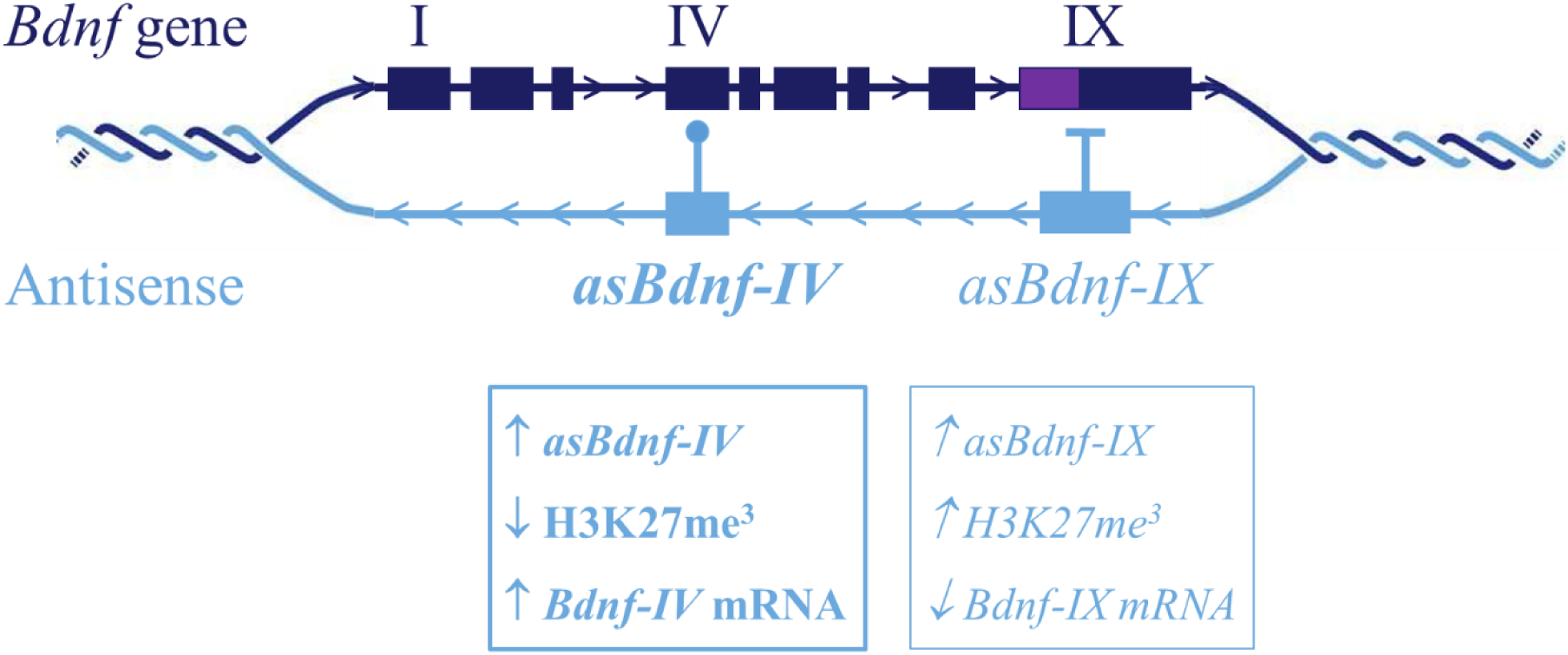
Summary of findings regarding asBdnf-IV regulation of Bdnf mRNA. The Bdnf gene has 9 exons, including exon IX that is common to all splice variants and contains the protein coding region of the gene (purple section). Here we described a novel antisense transcript opposite exon 4 of the Bdnf gene that is associated with a reduction in the repressive H3K27me^3^ (12) and increase in Bdnf-IV mRNA transcription. A second antisense transcript opposite exon 9 of the Bdnf gene (asBdnf-IX) has been previously reported (14) and exerts a repressive influence over Bdnf-IX gene expression through increased histone H3K27me^3^.

Importantly, following 6 days of extinction or abstinence from nicotine self-administration, changes in *asBdnf-IV* in the IL were not correlated with changes in *Bdnf* mRNA; and knockdown of the *asBdnf-IV* in the IL prior to extinction significantly attenuated reinstatement (for a previous report that BDNF regulates extinction of drug-seeking, see (8)). These findings suggest that the *asBdnf-IV* in the IL naturally *opposes* extinction learning, perhaps by maintaining or ‘priming’ an epigenetic state that is anathema to that extinction. We suspect that any such priming effect is highly specific to nicotine. This is supported by the ability of chronic nicotine to epigenetically prime the behavioural and molecular response to cocaine, whereas the inverse is not true (27). Moreover, within the PFC (including IL), *Bdnf* expression is known to be highly cell-specific (29) and nicotinic acetylcholine receptors (nAChRs) are widely expressed across layers I-IV on both glutamatergic projection neurons and GABAergic interneurons (30). Given that the epigenetic changes instigated by nicotine exposure are likely to occur via nAChR activation (28), and that many of these same cells express BDNF (31), it seems likely that the epigenetic regulation of BDNF in the IL following nicotine is strongly influenced by concurrent nAChR activation.

Given the increasing focus on lncRNA in human neurological and neurodegenerative disorders, the results of this study provide further evidence that chronic drug exposure significantly alters lncRNA expression. They are consistent with past reports that have linked *asBdnf-IX* expression with early-onset alcohol use disorder (32). We now extend this to show that, within the IL, *asBdnf-IV* plays a causative role in the extinction of nicotine-seeking. Taken together, this body of work confirms that BDNF signaling in the IL may be a useful target in the treatment of drug addiction, including development of anti-smoking medications.

## ACKNOWLEDGEMENTS

This work was supported by UNSW Faculty of Science research funding to KJC. K.J.C., N.A.Y. and K.V.M. conceived the study. K.J.C. and N.A.Y. designed the experiments, primers and oligonucleotides. K.J.C., M.C., A.S., K.K. and N.M.H. carried out the experiments. P.D.W. analysed sequencing data. K.J.C. performed statistical analysis. K.J.C. and N.M.H. wrote the manuscript.

